# Extraction, Characterization, and Toxicological Assessment of Chemicals from Thermal Bill Paper: An In-Silico ADME and Daphnia pulex Study

**DOI:** 10.1101/2024.08.27.609942

**Authors:** Usha Ramesh, Aniketh Vinod Bhat, Nalini Ranganath, Asha Changuli Hosamane, Athavan Alias Anand Selvam

**Affiliations:** Sai Krushna Vidya Mandir, Bengaluru, Karnataka, India; Rashtrotthana Vidya Kendra, Banashankari, Karnataka, India; Department of Chemistry, Prayoga Institute of Education Research, Bengaluru, Karnataka, India

**Keywords:** Chemical Extraction, Thermal Bill Paper, ADME properties, Daphnia Pulex, Bisphenol S

## Abstract

The current study provides a comprehensive analysis of the chemical composition of thermal bill paper, revealing the presence of hazardous chemical compounds such as Bisphenol S (BPS) and Diphenyl Sulfone (DPS). The extraction process using ethyl acetate as the solvent effectively isolated these compounds, as evidenced by Thin Layer Chromatography (TLC) results, which displayed distinct spots corresponding to the compounds of interest. Fourier Transform Infrared (FTIR) spectroscopy confirmed the presence of characteristic functional groups, while High-Resolution Mass Spectrometry (HR-MS) identified molecular weights consistent with BPS and DPS. In-silico analysis using the SwissADME web tool provided insights into the pharmacokinetic properties of these compounds, with bioavailability radar plots indicating optimal absorption ranges, except for a low degree of saturation. The BOILED-Egg model predicted that BPS is likely to be absorbed by the gastrointestinal tract, while DPS may cross the blood-brain barrier, raising potential concerns about its impact on the central nervous system. Toxicity studies conducted with Daphnia pulex showed significant toxicity at higher concentrations, supporting the idea that the presence of BPS and DPS in thermal bill paper poses a risk to human health and the environment. These findings underscore the need for increased regulation and the exploration of safer alternatives.

## 1. Introduction

Thermal paper is integral to modern transaction systems, particularly in receipt printing across sectors such as retail, banking, and hospitality. This specialized paper is coated with a chemical that changes colour when exposed to heat, eliminating the need for traditional ink-based printing methods [1]. Its efficiency and cost-effectiveness have made it a ubiquitous component in point-of-sale (POS) systems worldwide [2], streamlining transaction processes and enhancing customer service in various industries, including supermarkets, petrol stations, and healthcare. Figure 1 explains the mechanism of printing in the reactive layer of the thermal bill paper which includes a chemical called bisphenol A (BPA), a leuco dye, binders, and stabilizers [3]. In medical settings, thermal paper is critical for printing precise documents like medical test results, radiology images, and medication labels.

**Figure 1.**
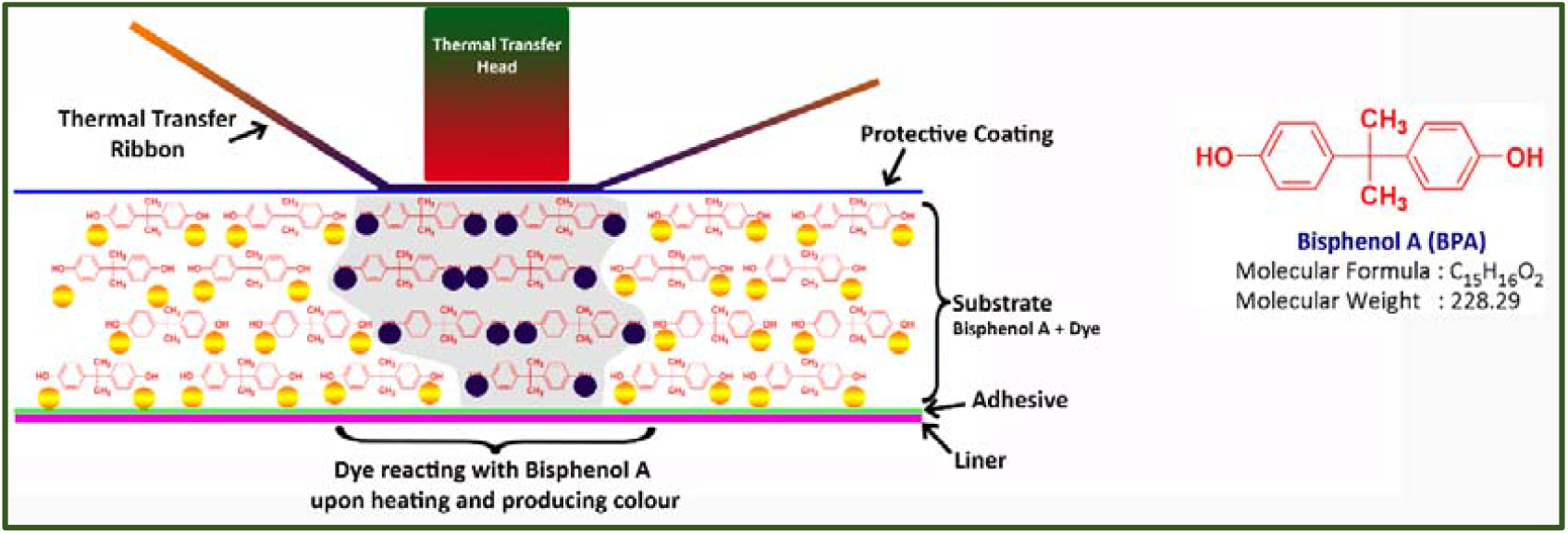
Mechanism of printing in the reactive layer of the thermal bill paper

Concerns regarding chemical hazards associated with thermal paper have been rising, particularly due to the presence of chemicals like Bisphenol A (BPA) and Bisphenol S (BPS) in its coating [4]. Beyond professional use, thermal paper is also common in everyday items such as cinema tickets, transport tickets, ATM receipts, and fax paper. Given that thermal paper is frequently handled in everyday transactions, the potential for chemical exposure and its implications on human health warrant serious attention and investigation. Bisphenol A (BPA) can function as a weak estrogenic receptor agonist and has been linked to numerous adverse health outcomes [5,6]. Exposure to BPA occurs not only through the food chain but also via inhalation from sources like floor dust and paints [7-9], as well as through skin contact (transdermal route). Thermal paper is a significant source of BPA exposure through the skin; studies have shown that 27% of BPA on the skin surface can penetrate and enter the bloodstream within two hours [10]. One of the primary health concerns associated with BPA is its endocrine-disrupting properties, which have been detected in human breast milk, blood, and urine samples [11-12]. BPA exposure has also been linked to issues such as impaired cognitive development, blood sugar irregularities, weight gain, and severe reproductive disorders [13]. The health risks associated with BPA have led to regulatory bans in countries such as the USA, Canada, and the European Union, prompting the development of alternative bisphenol compounds. While BPA exposure can occur through multiple routes due to its leaching from plastics and other products [14], dermal transfer has been specifically studied in humans who frequently handle thermal bill papers [15]. Later, a few derivatives of bisphenols such as bisphenol F (BPF), bisphenol S (BPS), bisphenol P (BPP), bisphenol B (BPB), and bisphenol AF (BPAF) were introduced in the market as an alternative to producing BPA-free products [16]. This shift has altered the profile of bisphenols (BPs) found in environmental and biological samples [17]. The structures of bisphenol analogues are shown in Figure 2. While these alternatives were introduced to mitigate the adverse effects associated with BPA, emerging studies suggest that they may also possess endocrine-disrupting properties. Understanding the implications of these chemicals is crucial for public health and environmental safety. The process of extracting and characterizing hazardous substances from thermal paper is vital in assessing their potential impact. We extracted chemicals from the thermal bill paper and characterised using appropriate solvents. Furthermore, the chemical structures were subjected to the Absorption, Distribution, Metabolism, and Excretion (ADME) properties prediction using Swiss-ADME online portal (www.swissadme.ch) for understanding their behaviour in biological systems and evaluating their potential toxicity [18,19].

**Figure 2.**
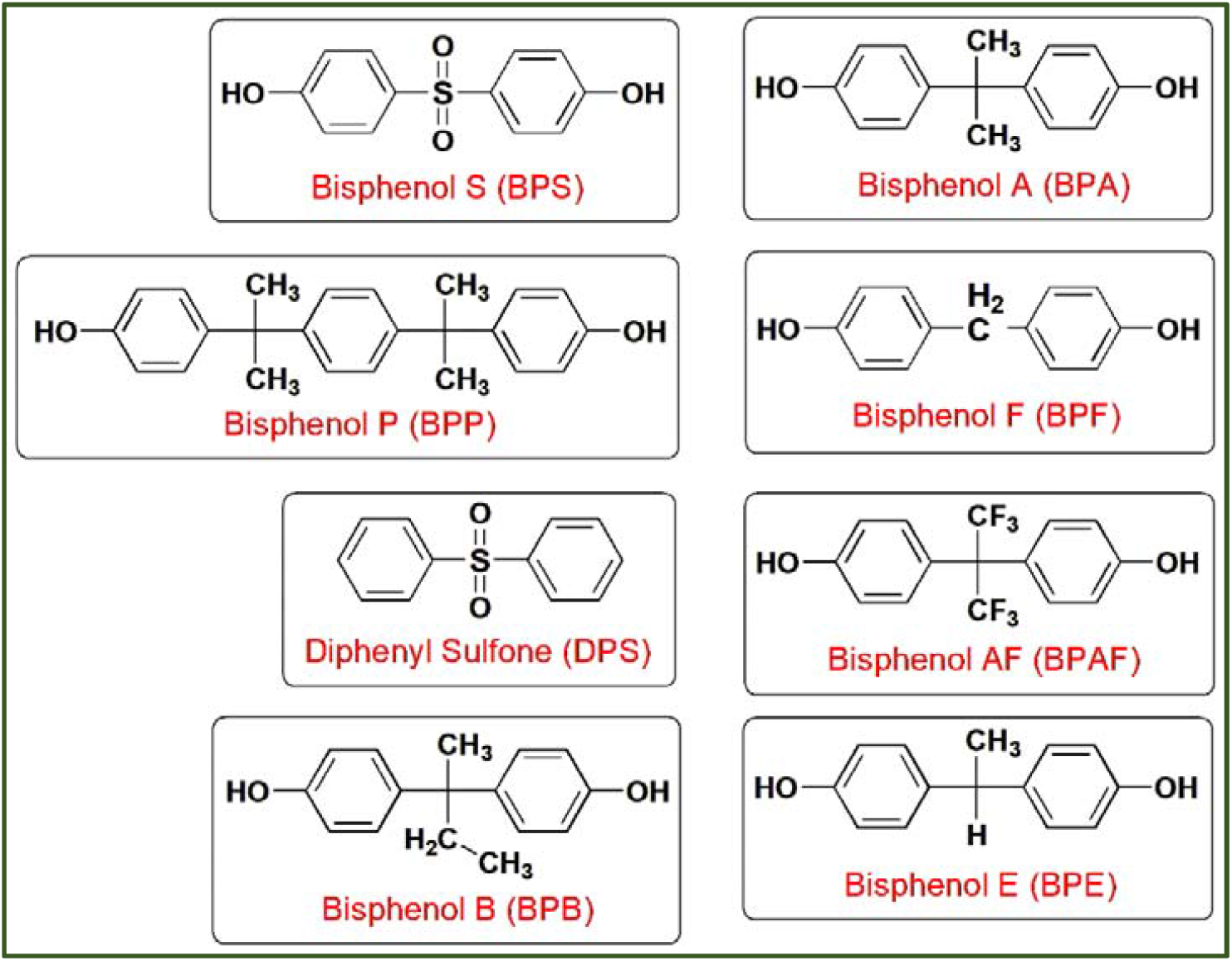
Derivatives of bisphenols present in plasticware and thermal bill paper

This study also focuses on toxicity assessments using Daphnia pulex, a freshwater crustacean widely used in ecotoxicological studies due to its sensitivity to environmental pollutants [20]. By evaluating the effects of thermal paper-extracted chemicals on Daphnia, this research aims to provide insights into the ecological risks posed by these substances, supporting the development of safer alternatives and informing regulatory policies [21]. The goals of this project are multi-faceted: extracting chemicals from thermal paper using appropriate solvents, characterizing them through FTIR and HRMS, predicting their ADME properties, and conducting toxicity studies using Daphnia pulex. This comprehensive approach aligns with the principles of sustainable chemistry, which aim to minimize the use and generation of harmful substances in chemical processes. The study contributes to global sustainability efforts by promoting responsible chemical management and reducing harmful chemical exposure.

## 2. Materials and Methods

### 2.1. Chemical Extraction from Thermal Bill Paper

All solvents and reagents were of analytical grade (purity>99%) and purchased from Sisco research lab products. The chemical extraction process for this study involved several key steps, focusing on the selection of solvents and extraction techniques to isolate compounds from thermal paper. Initially, unprinted thermal paper was purchased from a local stationary store, Bengaluru, India. The paper was cut into uniform 1×1 cm squares to ensure consistency in the extraction process. Various solvents such as hexane, petroleum ether, ethanol, chloroform, ethyl acetate, methanol, and water were tested for their efficacy in extracting compounds, with thin-layer chromatography (TLC). These experiments were used to determine the solvent that produced the most distinct spots, indicating the successful extraction of diverse compounds. To extract the chemicals, 5 grams of the cut thermal paper was subjected to extraction using different solvents. The extraction process was conducted using a water bath sonicator set at 60 °C for 30 minutes to enhance the efficiency of the solvent in penetrating and dissolving the paper’s coating. Following extraction, the solution was filtered and treated with a rotary evaporator to remove the ethyl acetate and concentrate the crude extract. The concentrated extract was then analyzed using TLC, revealing multiple spots, which suggested the presence of more than two compounds. For further characterization, the crude extract was analysed using high-resolution mass spectrometry (HRMS) and Fourier-transform infrared spectroscopy (FTIR) analysis.

### 2.2. Computational Study for ADME Properties Prediction

To predict the Absorption, Distribution, Metabolism, and Excretion (ADME) properties of the identified compounds, Swiss-ADME online portal was used. The chemical structures of the compounds, determined from the FTIR and MS analyses, were input into online predictive tools to assess their ADME characteristics. This tool evaluates several key ADME parameters, including physicochemical properties, lipophilicity, water solubility, pharmacokinetics, druglikeness, and medicinal chemistry properties.

The five steps for pharmacophore modelling are given below:

#### 2.2.1 Physicochemical Properties

SwissADME’s physicochemical properties section provides predictions for molecular attributes affecting the pharmacokinetics and pharmacodynamics of small molecules. Key properties include molecular weight, logP (lipophilicity), number of atomic heavy atoms, hydrogen bond donors and acceptors, refractivity, fraction Csp3, TPSA (topological polar surface area), and the number of rotatable bonds. These properties serve as crucial descriptors in models and rules for quickly estimating significant ADME properties like absorption and brain access.

#### 2.2.2 Lipophilicity

The ability of compounds to cross biological barriers and reach specific targets is influenced by their lipophilicity. Highly lipophilic compounds exhibit better membrane permeability and can cross biological barriers more easily, but they also tend to bind non-specifically and accumulate in lipid-rich tissues. On the other hand, compounds with low lipophilicity show lower bioavailability and limited access to certain targets but are less likely to cause off-target effects or toxicity. SwissADME provides five predictive models for lipophilicity, including XLOGP3 (an atomistic method using corrective factors), WLOGP (a fragment-based atomistic method), MLOGP (a topological method using 13 molecular descriptors), SILICOS-IT (a hybrid method utilizing 27 fragments and 7 topological descriptors), and iLOGP (a physics-based method employing the Generalized-Born and solvent accessible surface area (GB/SA) model). The platform also offers a consensus log Po/w value, averaging the predictions from all five methods.

#### 2.2.3 Water Solubility

Water solubility is a critical factor in drug discovery as it impacts a compound’s bioavailability and pharmacokinetic properties. SwissADME predicts water solubility using three models: ESOL (Estimated SOLubility), a model by Ali et al., and SILICOS-IT, a machine learning-based approach. These models are trained on extensive datasets of experimentally measured solubility values, providing predictions as the decimal logarithm of molar solubility in water (log S). SwissADME also offers solubility values in mol/L and mg/mL, along with qualitative solubility classes.

#### 2.2.4 Pharmacokinetics

In this section, SwissADME evaluates specific ADME behaviors of the organic compound in question. Predictions for passive human gastrointestinal absorption (HIA) and blood-brain barrier (BBB) permeation are presented using the BOILED-Egg model, a straightforward graphical classification method. The assessment includes whether compounds are substrates or non-substrates of the permeability glycoprotein (P-gp), which is essential for evaluating active efflux through biological membranes, such as from the gastrointestinal wall to the lumen or the brain. Additionally, the platform predicts inhibition of five major cytochrome P450 enzymes (CYP1A2, CYP2C19, CYP2C9, CYP2D6, CYP3A4) and the skin penetration coefficient (Log Kp). These predictions are instrumental in optimizing pharmacokinetics and evaluating small drug-like organic compounds.

#### 2.2.5 Druglikeness

SwissADME predicts druglikeness based on five rule-based filters: Lipinski, Ghose, Veber, Egan, and Muegge. The criteria are:

- Lipinski’s rule: Molecular weight ≤ 500, MLOGP (lipophilicity) ≤ 4.15, hydrogen bond acceptors ≤ 10, hydrogen bond donors ≤ 5.
- Ghose’s rule: Molecular weight between 160 and 480, WLOGP (lipophilicity) between -0.4 and 5.6, molar refractivity between 40 and 130, number of atoms between 20 and 70.
- Veber’s rule: Number of rotatable bonds ≤ 10, total polar surface area ≤ 140.
- Egan’s rule: WLOGP (lipophilicity) ≤ 5.88, total polar surface area ≤ 131.6.
- Muegge’s rule: Molecular weight between 200 and 600, XLOGP3 (lipophilicity) between -2 and 5, total polar surface area ≤ 150, number of rings ≤ 7, number of carbon atoms > 4, number of heteroatoms > 1, number of rotatable bonds ≤ 15, hydrogen bond acceptors ≤ 10, hydrogen bond donors ≤ 5.

#### 2.2.6 Medicinal Chemistry

This section aims to assist medicinal chemists in drug discovery efforts. It includes four parameters: PAINS (pan assay interference compounds): Identifies compounds with substructures that show potent responses in assays regardless of the protein target. Brenk: Provides information on potentially toxic, chemically reactive, metabolically unstable compounds, or those with poor pharmacokinetic properties. Leadlikeness: Similar to drug-likeness, focusing on physicochemical boundaries defines a good lead compound. Synthetic Accessibility (SA): Summarizes fragmental contributions to SA, corrected by terms describing size and complexity, such as macrocycles, chiral centers, or spiro functions. The SA score helps prioritize molecules for synthesis.

### 2.3 Toxicity Assessment using Daphnia Pulex

Daphnia cultures were maintained following the method described by Stark and Vargas (2003) [22]. They were fed a solution containing a 1:1 mixture of yeast–cereal leaves–trout chow (YCT) and the algal species *Selenastrum capricornutum*. The Daphnia were reared in reconstituted dilution water (RDW) within a freestanding environmental chamber set to 25°C, 50% relative humidity, and a 16-hour light/8-hour dark cycle. To evaluate the potential health and environmental risks of the extracted chemicals, toxicity studies were conducted using the freshwater crustacean, Daphnia pulex, which was purchased from Microworms Company. Daphnia pulex, commonly known as water fleas, are widely used in ecotoxicology due to their sensitivity to pollutants and their importance in aquatic ecosystems.

The crude chemical extract was dissolved in water to create solutions at microgram concentration levels. Various concentrations such as 0.25, 0.5, 1.0, 2.0 and 5.0 mg/L of the crude product were prepared to assess dose-dependent effects. Equal numbers of Daphnia were placed in 5 mL beakers containing the different concentrations of the crude extract solution. Control groups were maintained in water without the extract to compare the effects. The Daphnia were exposed to the chemical solutions, and their responses were monitored over different time periods, 2, 4, 6, 24, 48 and 72 hours. Mortality was recorded at both time points to determine the lethal concentration of the extract.

#### 2.3.1 Heartbeat Rate

After different time periods of exposure, the heartbeat rates of the Daphnia were monitored. The heartbeat rate is a sensitive indicator of physiological stress and toxicity in Daphnia, providing insights into the sub-lethal effects of chemical exposure. The data from these toxicity studies, including survival and heartbeat rates, helped to evaluate the acute toxicity of the extracted compounds. By assessing both lethal and sub-lethal endpoints, the study provided a comprehensive understanding of the potential hazards posed by the chemicals present in thermal paper. These findings are critical for informing risk assessments and guiding the development of safer alternatives.

## 3 Results and Discussion

### 3.1 Chemical Extraction from Unprinted Thermal Bill Paper Roll

Various solvents were employed to extract compounds from the thermal bill roll paper. Among the solvents tested, ethyl acetate was identified as the most effective solvent based on the results of TLC. The TLC analysis revealed three prominent spots, likely representing two chemicals for coating and one dye compound. The mobile phase used for the TLC was a 1:1 mixture of ethyl acetate and hexane. The TLC plate was examined under UV light to visualize the spots (Figure 3). The crude extract, obtained after solvent evaporation, appeared black and was subsequently characterized using Fourier-transform infrared spectroscopy (FTIR) and high-resolution mass spectrometry (HRMS). From 5 grams of thermal bill paper, approximately 0.510 grams of chemical compounds were extracted.

**Figure 3.**
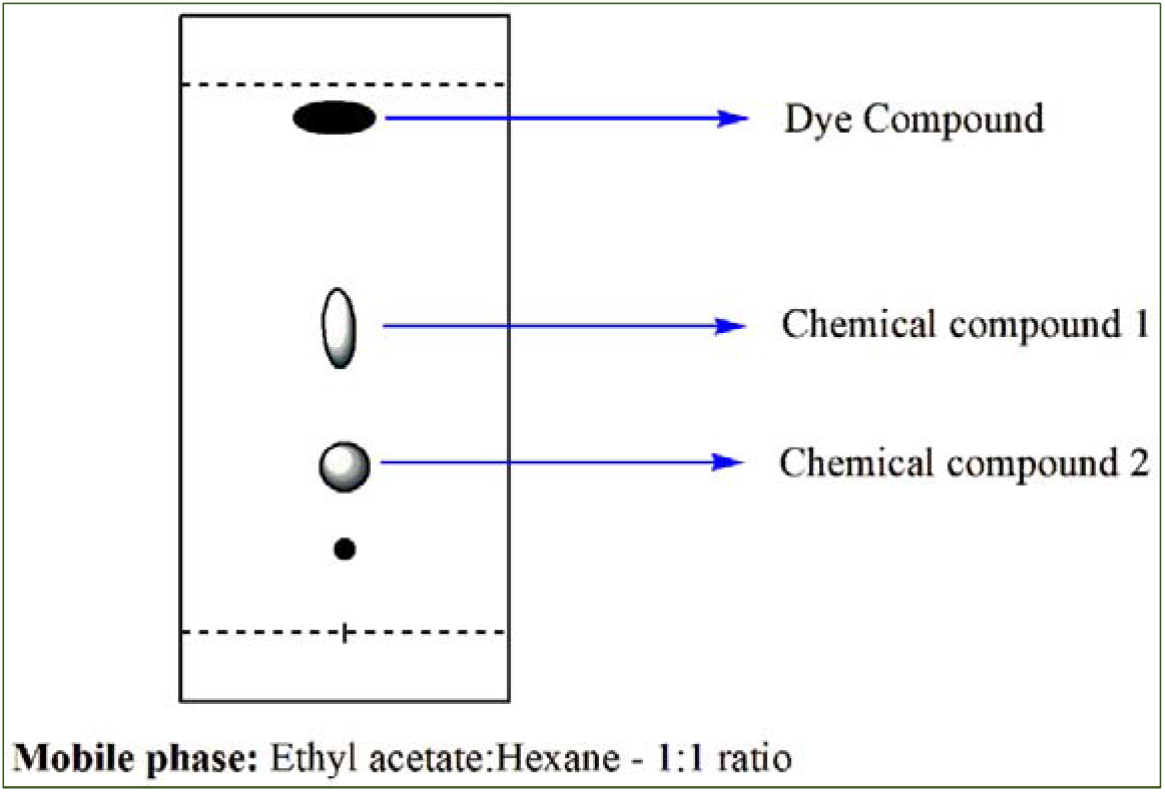
Thin layer chromatography (TLC) plate with compounds eluted using an ethyl acetate-hexane mixture

### 3.2 FTIR Analysis

The FTIR analysis of the crude compound was performed using the ATR technique. The FTIR spectral data are given in Table 1. The information on the functional groups present in the compound was identified using the FTIR peaks. The FTIR spectrum analysis of the chemical extracts from the thermal bill paper reveals distinct absorption bands corresponding to specific functional groups, indicating the presence of certain bisphenol compounds. A broad absorption band at 3346 cm◻^1^ is observed, which can be attributed to O-H symmetric stretching, suggesting the presence of hydroxyl groups. The peaks at 2916 cm◻^1^ and 2849 cm◻^1^ are characteristic of aromatic C-H stretching, further supporting the presence of aromatic compounds. The band at 1638 cm◻^1^ corresponds to C=C stretching, commonly associated with the aromatic ring structures found in bisphenols. Additionally, the absorption at 1290 cm◻^1^ is indicative of C-O symmetric stretching, while the peak at 1148 cm◻^1^ can be assigned to S=O stretching. Together, these spectral features confirm the presence of bisphenol-related compounds in the extracts, aligning with the known chemical composition of thermal bill paper.

**Table 1.**
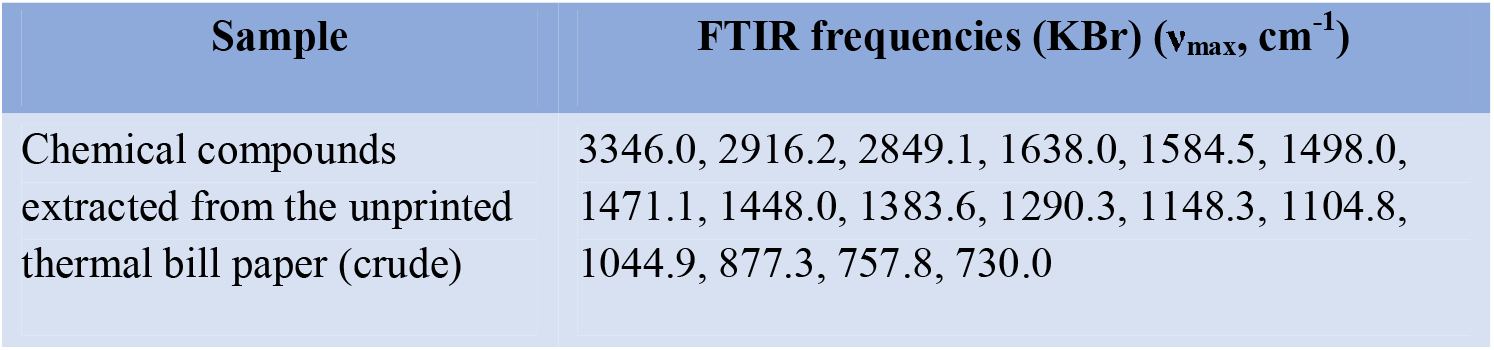
FTIR spectral data of the chemical extracted from the thermal bill paper.

| Sample | FTIR frequencies (KBr) ($\nu_{\text{max}}$, cm<sup>-1</sup>) |
| --- | --- |
| Chemical compounds extracted from the unprinted thermal bill paper (crude) | 3346.0, 2916.2, 2849.1, 1638.0, 1584.5, 1498.0, 1471.1, 1448.0, 1383.6, 1290.3, 1148.3, 1104.8, 1044.9, 877.3, 757.8, 730.0 |

**Table 2.** The output table by the SwissADME tool for BPS and DPS represents its drug-like properties.

|  | Compound Name | Bisphenol S (BPS) | Diphenyl Sulfone (DPS) |
| --- | --- | --- | --- |
|  | Canonical SMILES | OC1=CC=C(C=C1)S(=O)(=O)C1=CC=C(O)C=C1 | O=S(=O)(C1=CC=CC=C1)C1=CC=CC=C1 |
| **(i) Physiochemical Properties** |  |  |  |
| 1 | Formula | C<sub>12</sub>H<sub>10</sub>O<sub>4</sub>S | C<sub>12</sub>H<sub>10</sub>O<sub>2</sub>S |
| 2 | Molecular Weight | 250.27 | 218.27 |
| 3 | Number of heavy atoms | 17 | 15 |
| 4 | Number of aromatic heavy atoms | 12 | 12 |
| 5 | Fraction Csp<sup>3</sup> | 0 | 0 |
| 6 | Number of rotatable bonds | 2 | 2 |
| 7 | Number of H-bond acceptors | 4 | 2 |
| 8 | Number of H-bond donors | 2 | 0 |
| 9 | Topological polar surface area (TPSA) | 82.98 | 42.52 |
| **(ii) Lipophilicity** |  |  |  |
| 1 | Consensus Log $P_{o/w}$ | 1.88 | 2.68 |
| **(iii) Water Solubility** |  |  |  |
| 1 | Silicos-IT LogSw | -3.69 | -4.85 |
| **(iv) Pharmacokinetics** |  |  |  |
| 1 | GI absorption | High | High |
| 2 | BBB permeant | No | Yes |
| 4 | CYP1A2 inhibitor | No | No |
| 5 | CYP2C19 inhibitor | No | No |
| 6 | CYP2C9 inhibitor | No | No |
| 8 | CYP3A4 inhibitor | No | No |
| **(v) Drug likeness** |  |  |  |
| 1 | Lipinski number of violations | Yes, 0 violation | Yes, 0 violation |
| 2 | Ghose number of violations | Yes | Yes |
| 3 | Veber number of violations | Yes | Yes |
| 4 | Egan number of violations | Yes | Yes |
| 5 | Muegge number of violations | Yes | Yes |
| **(vi) Medicinal Chemistry** |  |  |  |
| 1 | PAINS number of alerts | 0 alert | 0 alert |
| 2 | Brenk number of alerts | 0 alert | 0 alert |
| 3 | Lead likeness | Yes | No; 1 violation: MW<250 |
| 4 | Synthetic accessibility | 1.88 | 1.90 |

**Table 3.** Toxicity study using *Daphnia pulex* at various concentrations and time intervals.

| Concentration of chemicals mixture | Time of Exposure (hours) |  |  |  |  |  |
| --- | --- | --- | --- | --- | --- | --- |
|  | 2 | 4 | 6 | 24 | 48 | 72 |
| Control | 0 | 0 | 0 | 8 | 15 | 20 |
| 0.25 mg/L | 0 | 0 | 0 | 20 | 35 | 54 |
| 0.5 mg/L | 0 | 0 | 10 | 25 | 35 | 55 |
| 1 mg/L | 0 | 15 | 15 | 40 | 50 | 65 |
| 2 mg/L | 0 | 20 | 20 | 40 | 60 | 65 |
| 5 mg/L | 40 | 40 | 60 | 100 | 100 | 100 |

### 3.3 HR-MS Analysis

The results of the HR-MS spectrum of the chemical extracts from the thermal bill paper reveal two prominent peaks at 251.0390 and 219.0485 m/z, which provide crucial insights into the molecular composition of the sample. By correlating the HR-MS peaks with known molecular weights, and supported by a thorough literature survey, it can be justified that the two major compounds identified in the thermal bill paper are Bisphenol S (BPS) and Diphenyl Sulfone (DPS) (Figure 4). Bisphenol S, with a molecular weight of 250, corresponds with the HR-MS peak at 251.0390 m/z, indicating its presence in the sample. The peak at 219.0485 m/z aligns with Diphenyl Sulfone, which has a molecular weight of 218. This alignment of molecular weights with the observed HR-MS peaks, supported by FTIR analysis that identifies functional groups consistent with these compounds, strongly justifies the conclusion that Bisphenol S and Diphenyl Sulfone are the primary constituents in the chemical extracts from the thermal bill paper. These findings are consistent with known compositions of thermal paper, which often include these compounds due to their functional properties in thermal printing applications.

**Figure 4.**
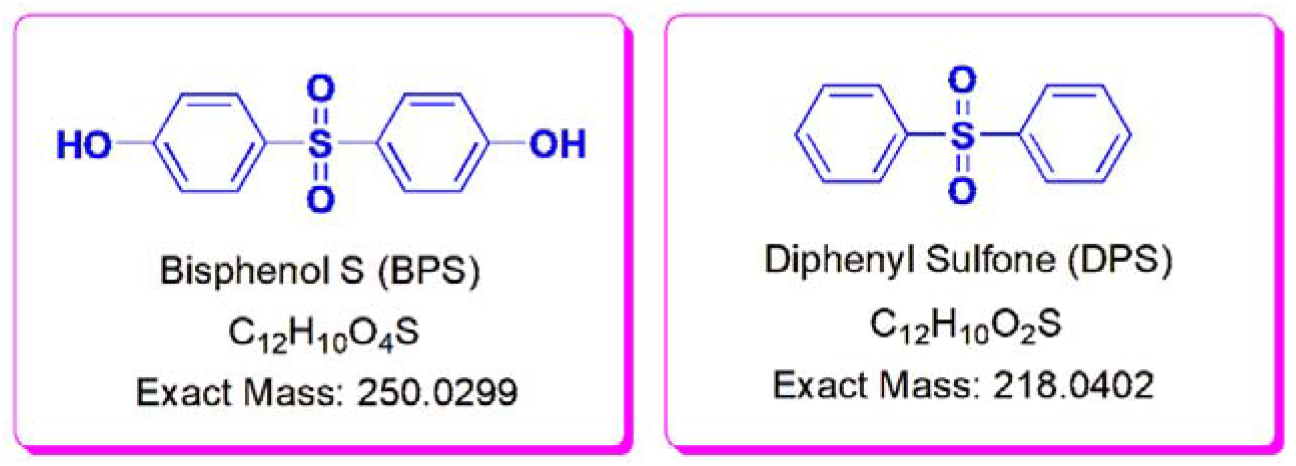
Chemical compounds extracted and confirmed by FTIR and HRMS

### 3.4 ADME Properties Prediction using SwissADME Web Tool

The in-silico drug-likeness properties of the BPS and DPS chemical structures were analyzed using the SwissADME online tool. The predictions and analysis were based on the numerical data obtained under the headings (i) physiochemical properties, (ii) lipophilicity, (iii) water solubility, (iv) pharmacokinetics, and (v) medicinal chemistry. The results included a bioavailability radar which represents six physicochemical properties of BPS and DPS: (i) Lipophilicity (XLOGP3 between −0.7 and +5.0), (ii) size (molecular weight between 150 and 500 g/mol), (iii) polarity (the total polar surface area between 20 and 130 Å2), (iv) solubility (log S not higher than 6), (v) saturation (fraction Csp3 not less than 0.25), and (vi) flexibility (the number of rotatable bonds not more than 9).

The bioavailability radars of BPS and DPS, upon interaction for six physicochemical properties, are shown in Figure 5. The efficiency of the BOILED-Egg method in predicting human blood-brain barrier (BBB) penetration and gastrointestinal absorption was also demonstrated. The BOILED-Egg representations of BPS and DPS created by SwissADME are displayed in Figure 6.

**Figure 5.**
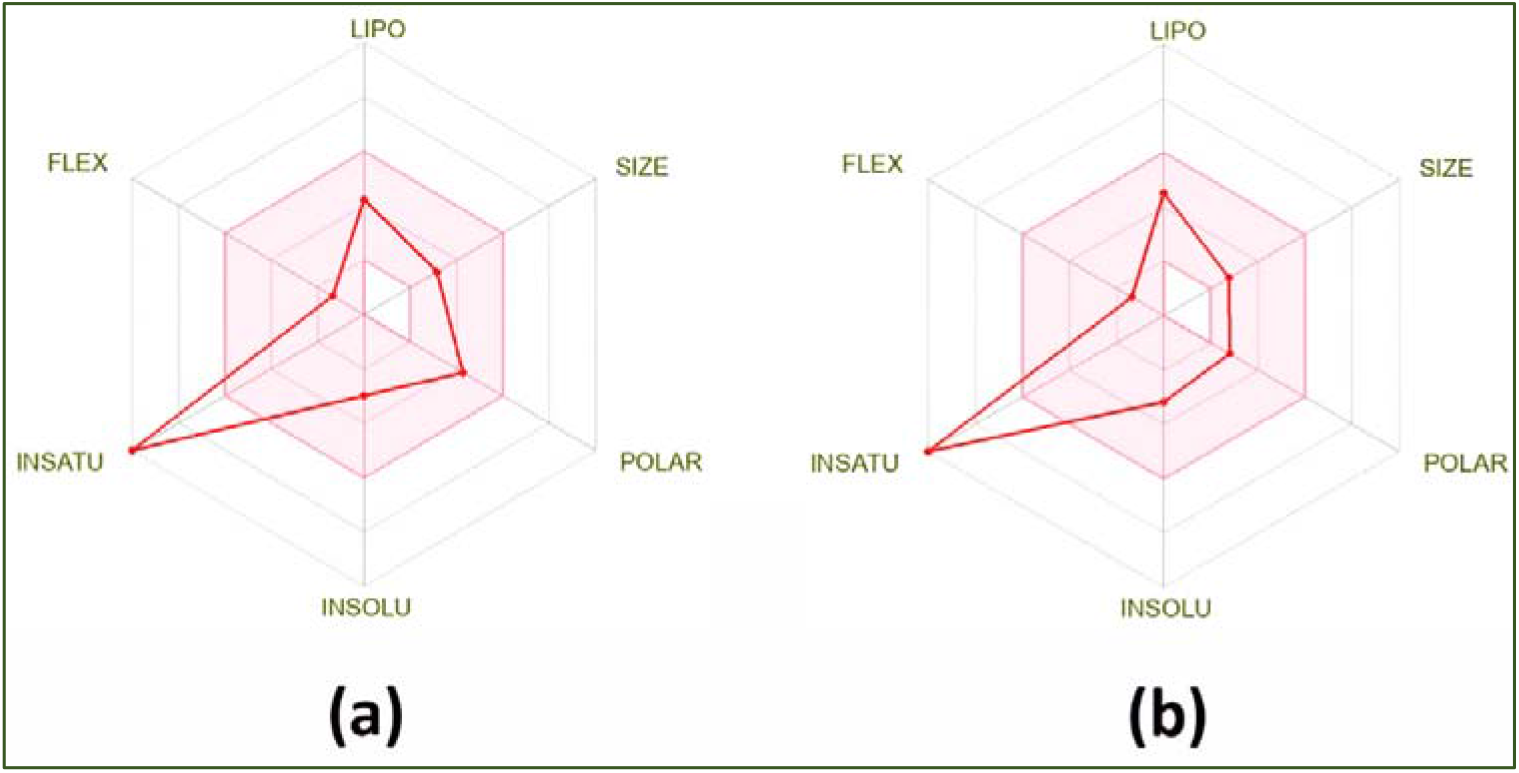
Bioavailability radar plot for (a) BPS and (b) DPS by SwissADME web tool

**Figure 6.**
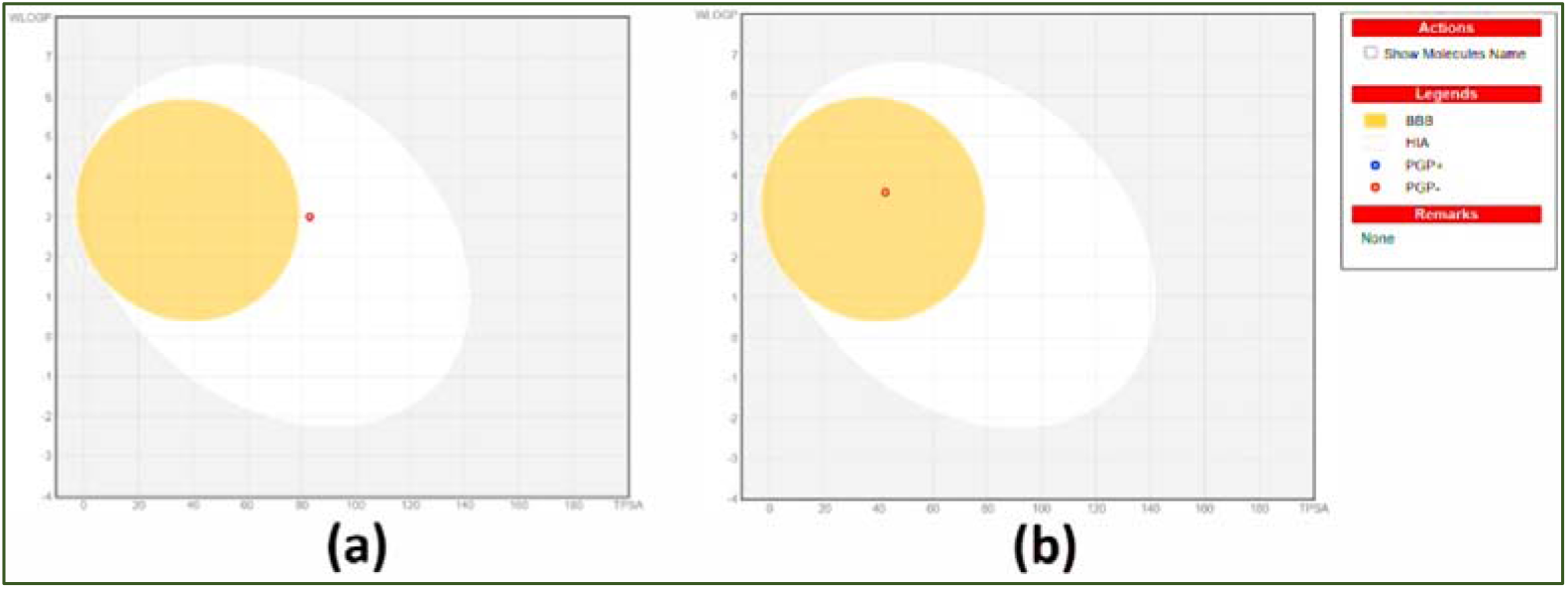
The BOILED-Egg prediction generated by SwissADME for (a) BPS and (b) DPS

In the bioavailability radar of BPS and DPS, the pink area represented the optimal range of these properties, and the red line represented the properties of the compounds. The bioavailability radar figure showed that the red lines of the BPS and DPS were in the range of the pink area except for INSATU i.e., about the saturation of BPS and DPS. The data showed the value for Csp3 was 0 for both compounds. Since the values for BPS and DPS were 0 (<0.25), the INSATU values in the radar plot were not appreciated by the software. According to the software program, highly saturated compounds are more likely to give good results in clinical trials. Research reports also showed saturation compounds are highly soluble with low melting points [23]. According to the BOILED-Egg predictions generated by SwissADME for BPS and DPS, BPS was located in the white region with a red point. This indicates that the BPS was likely to be absorbed by the gastrointestinal tract. DPS was located in the yellow region with a red point and this indicated that DPS was likely to be absorbed by the gastrointestinal tract and also pass through the blood-brain barrier (BBB). Additionally, the red point signified that both BPS and DPS were PGP-, and not effluxed from the central nervous (CNS) system by the P-glycoprotein. P-glycoprotein is a type of transporter protein that plays a crucial role in the BBB. Using the Silicos IT LogSw descriptor of SwissADME, the water solubility of BPS and DPS was predicted and its LogSw values were determined to be −3.69 and -4.85. According to the SwissADME LogSw scale, compounds with values less than (more negative than) −6 were classified as poorly soluble.

The assessment of lipophilicity was conducted through the prediction of the logarithm of the n-octanol/water partition coefficient using the Consensus LogPo/w descriptor of SwissADME. To achieve good oral bioavailability, which entails good permeability and solubility, a moderate logP value between 0 and 3 was recommended [24]. The predicted values of LogPo/w for BPS and DPS were 1.88 and 2.68. SwissADME was employed to estimate the metabolism of the BPS and DPS by inhibiting the principal cytochromes (CYP) of the P450 superfamily, including CYP1A2, CYP2C19, CYP2C9, and CYP3A4. CYP enzyme inhibition, a primary mechanism for drug-drug interactions based on metabolism, involves competition with other drugs for the same enzyme binding site. The inhibition of enzymes impairs the biotransformation or clearance of clinically used drugs, leading to elevated plasma levels of drugs that affect the therapeutic outcome. In the case of a prodrug, the effect is reduced. Therefore, the inhibition of CYPs resulted in drug toxicity or lack of efficacy. The predicted outcomes suggested that both BPS and DPS were not inhibit cytochromes of the P450 superfamily. Finally, the structures of BPS and DPS underwent drug-likeness prediction using five different rule-based filters, namely Lipinski, Ghose, Veber, Egan, and Muegge. The evaluation showed that both the compounds complied with all rules and did not violate any of them, indicating good drug-likeness.

### 3.5 Toxicity Studies using *Daphnia Pulex*

Acute toxicity tests indicated that both the chemicals from thermal bill paper at higher concentrations had a detrimental effect on the survival of *Daphnia pulex*. The initial toxicity effects at 5 mg/L concentration were noticed after 2 hours of contact period, as depicted in Figure 7. As anticipated, the *Daphnia* death rate was lowest in the controls and increased, although not linearly, with increasing chemical concentration to 5 mg/L. The heart rate of *Daphnia pulex* serves as a valuable biomarker for assessing the cardiotoxic effects of chemical exposure. By studying the cardiac events in *Daphnia pulex* when exposed to chemicals extracted from thermal bill paper, researchers can gain insights into the potential toxic effects these substances may have on cardiac function. The need for studying the heartbeat rate in *Daphnia pulex* arises from its effectiveness as a biomarker for assessing the cardiotoxic effects of various chemicals, including those extracted from thermal bill paper.

**Figure 7.**
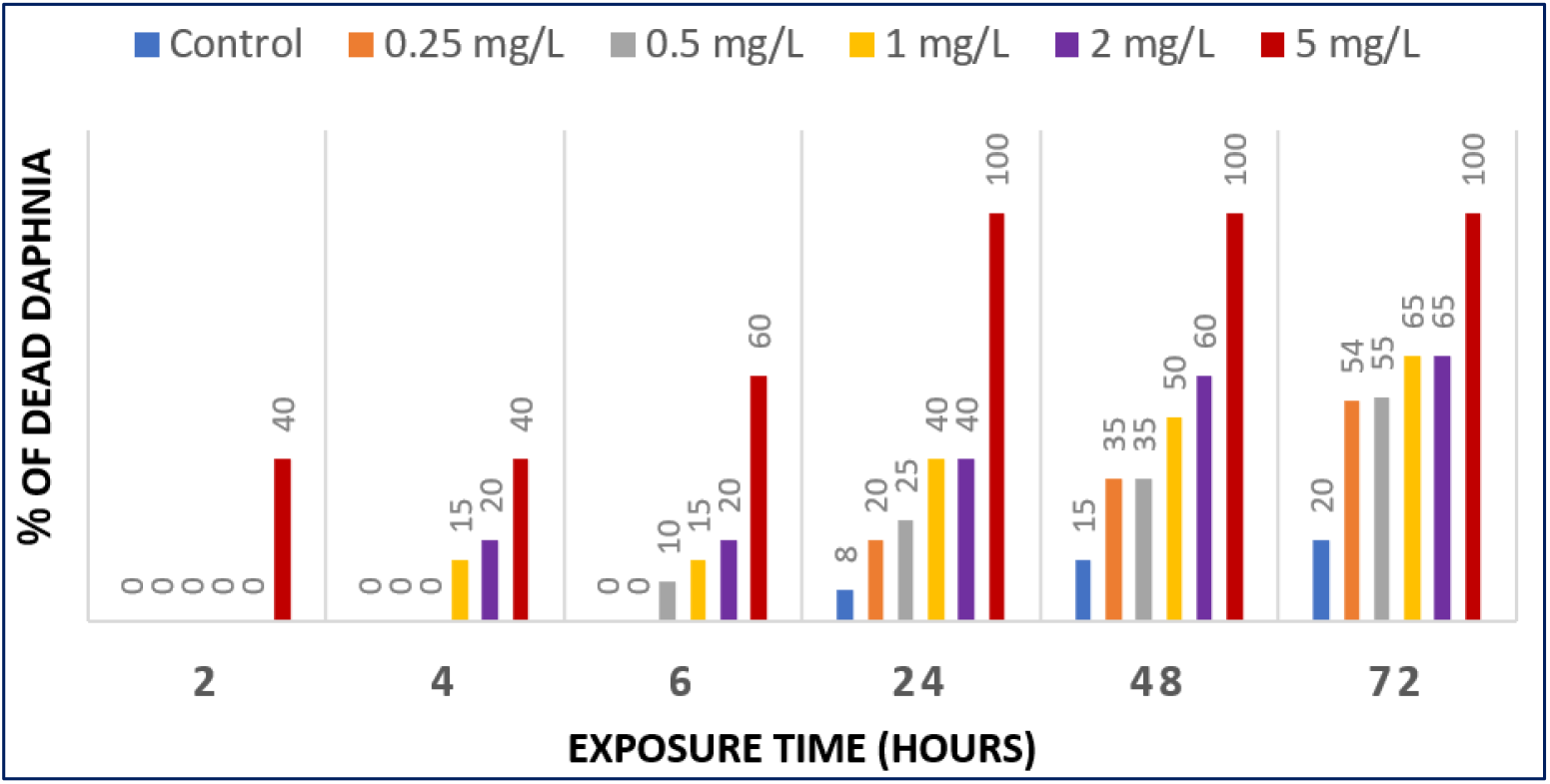
*Daphnia pulex* for toxicity assessment at various concentrations and time intervals

**Figure 8.**
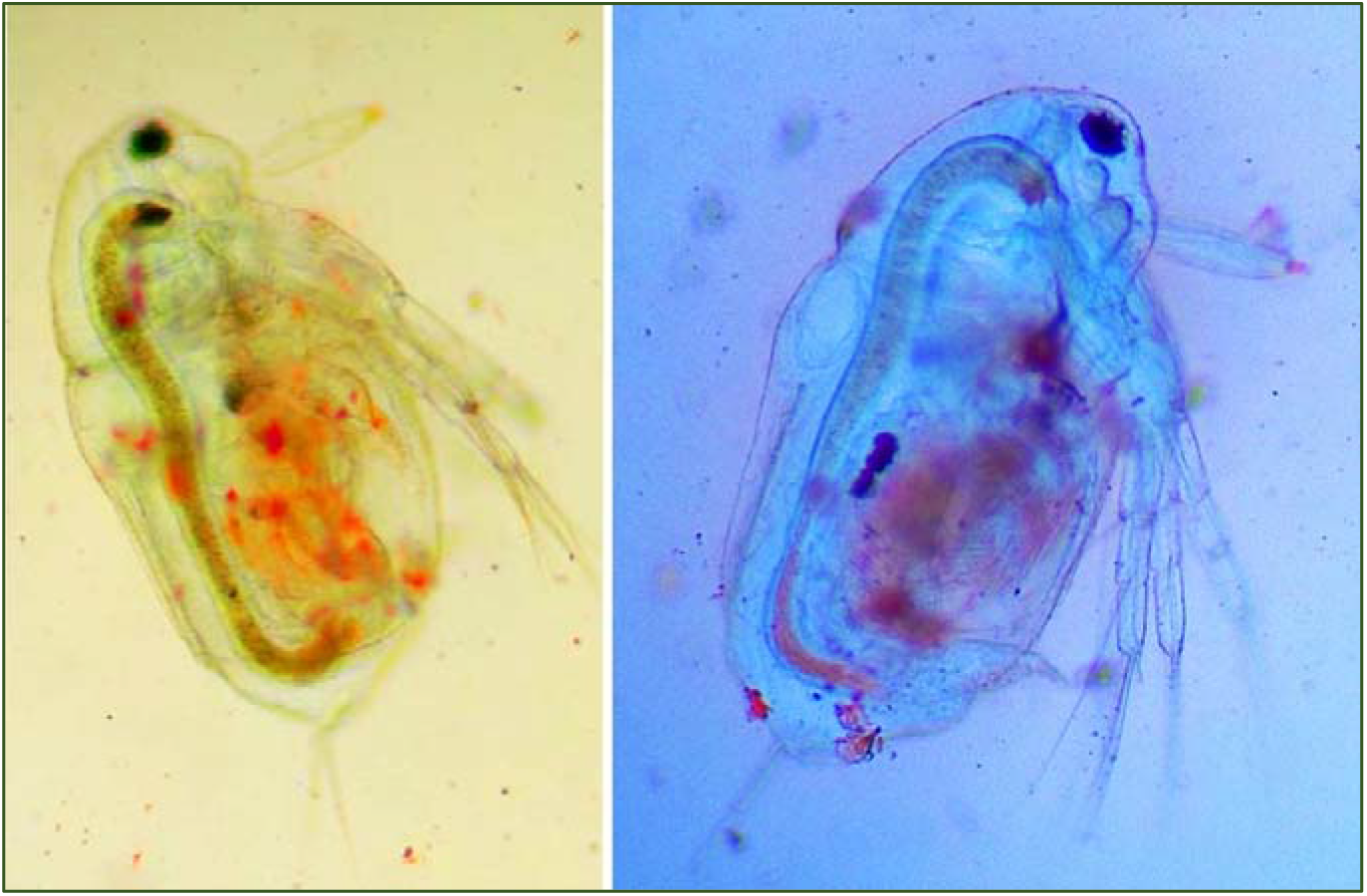
Microscopic analysis of *Daphnia pulex* for assessing heart beat rate

**Figure 9.**
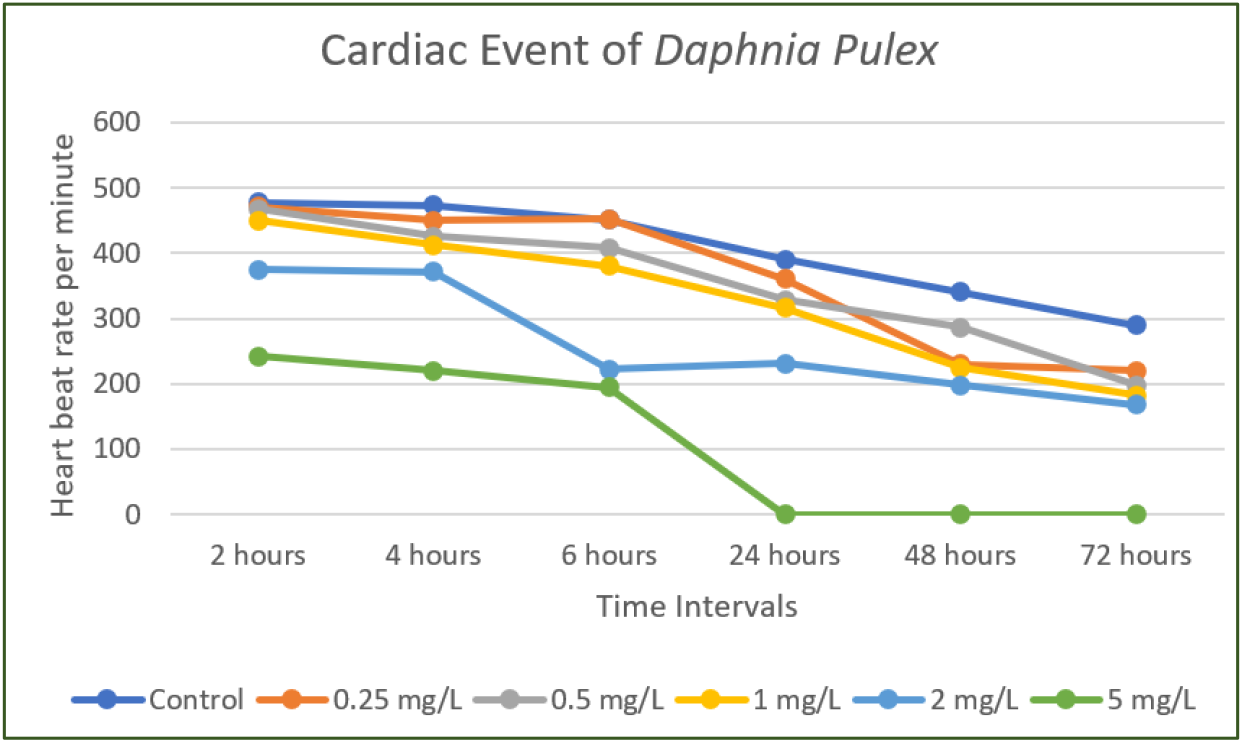
Trends in *Daphnia pulex* heart rate across various concentrations of chemicals from thermal bill paper

The heart of *Daphnia pulex* is highly sensitive to toxicants, making changes in its heart rate a direct and quantifiable indicator of stress and toxicity. Monitoring these changes is crucial in understanding the broader toxicological impacts of thermal bill paper chemicals, beyond specific environmental concerns. Unlike studies focused solely on aquatic ecosystems, this approach offers a non-specific screening method to assess the systemic effects of chemical exposure, providing insights into overall physiological distress. Moreover, since chemicals like Bisphenol S (BPS), found in thermal bill paper, are known to affect cardiovascular health in humans, studying their impact on *Daphnia pulex* can offer predictive value for potential human health risks. The use of *Daphnia pulex* allows for rapid, ethical testing, yielding immediate data on cardiotoxicity that can guide further toxicological evaluations and inform public health recommendations.

The results from the study show a clear dose-dependent decrease in the heart rate of *Daphnia pulex* when exposed to increasing concentrations of the chemical extracted from thermal bill paper over various time intervals. The cardiac event data for *Daphnia pulex* at various concentrations and time intervals are presented in Table 4. At the 2-hour mark, the heart rates of the *Daphnia* were relatively stable across all concentrations, with the control group showing a rate of 478 beats per minute (bpm) and the highest concentration (5 mg/L) showing a significant reduction to 242 bpm. As time progressed, the heart rates in all groups exposed to the chemical steadily declined.

**Table 4.** The cardiac event of *Daphnia pulex* recorded at various concentrations and time intervals.

|  | 2 hours | 4 hours | 6 hours | 24 hours | 48 hours | 72 hours |
| --- | --- | --- | --- | --- | --- | --- |
| **Control** | 478 | 473 | 451 | 390 | 340 | 290 |
| **0.25 mg/L** | 471 | 450 | 452 | 360 | 230 | 220 |
| **0.5 mg/L** | 468 | 426 | 408 | 328 | 286 | 198 |
| **1 mg/L** | 450 | 412 | 380 | 316 | 224 | 182 |
| **2 mg/L** | 375 | 372 | 222 | 231 | 198 | 168 |
| **5 mg/L** | 242 | 220 | 195 | 0 | 0 | 0 |

By the 24-hour mark, a more pronounced reduction in heart rate was observed, particularly in the groups exposed to higher concentrations. The heart rate in the 5 mg/L group dropped to 0 bpm, indicating complete cardiac arrest, while the control group maintained a heart rate of 390 bpm. The pattern continued at the 48-hour and 72-hour intervals, where the higher concentrations (1 mg/L, 2 mg/L, and 5 mg/L) showed dramatic decreases in heart rate, with the 5 mg/L group showing no heartbeats after 24 hours. In contrast, the control group experienced a gradual decline but maintained a heartbeat throughout the 72-hour period, ending at 290 bpm. These findings suggest that the chemical extracted from thermal bill paper has a significant cardiotoxic effect on *Daphnia pulex*, with the severity of the effect increasing both with higher concentrations and longer exposure times.

## 4. Conclusion

The study provided a comprehensive analysis of the chemical composition of thermal bill paper, revealing the presence of potentially hazardous compounds such as Bisphenol S (BPS) and Diphenyl Sulfone (DPS). The extraction process using ethyl acetate as the solvent was effective, as evidenced by the TLC results showing distinct spots corresponding to the compounds of interest. The subsequent FTIR and HR-MS analyses confirmed the presence of these compounds by identifying characteristic functional groups and matching molecular weights with known standards. The identification of BPS and DPS as major constituents aligns with their known usage in thermal paper due to their functional properties.

The in-silico analysis using the SwissADME web tool provided significant insights into the pharmacokinetic properties of BPS and DPS. The bioavailability radar plots indicated that both compounds possess properties within the optimal range for absorption and bioavailability, except for their low degree of saturation, which could affect their clinical efficacy. The BOILED-Egg model predicted that BPS is likely to be absorbed by the gastrointestinal tract, while DPS could potentially cross the blood-brain barrier, raising concerns about its impact on the central nervous system. The lack of P-glycoprotein recognition further implies that these compounds might persist in the CNS, potentially leading to adverse neurological effects. The toxicity studies conducted using Daphnia pulex provided additional evidence of the potential harm posed by these chemicals. The acute toxicity tests demonstrated that at higher concentrations, both BPS and DPS exhibit significant toxicity, further supporting the notion that their presence in thermal bill paper could pose a serious risk to human health and the environment. These findings are consistent with existing literature, which has raised concerns about the widespread use of bisphenol compounds in consumer products and their potential endocrine-disrupting effects.

In conclusion, the study highlights the need for increased scrutiny and regulation of the chemicals used in thermal bill paper. The presence of BPS and DPS, coupled with their potential to persist in human tissues and cause adverse health effects, underscores the importance of finding safer alternatives. The use of advanced analytical techniques such as FTIR and HR-MS, combined with in-silico tools like SwissADME, provides a robust framework for assessing the safety of chemical compounds in consumer products. As the understanding of the health risks associated with bisphenol compounds continues to evolve, it is crucial to explore and adopt safer alternatives to protect public health and the environment.

## Acknowledgements

We are thanking Indian Institute of Science for providing the library and instrumentation facilities.

## Declarations

### Ethical Approval

“not applicable”.

### Availability of data and materials

The data and materials are attached as supporting information.

