## Supplementary Material for "Extraction, Characterization, and Toxicological Assessment of Chemicals from Thermal Bill Paper: An In-Silico ADME and Daphnia pulex Study"

*Corresponding author

**Table S1** FTIR band assignments of the chemicals extracted from the thermal bill paper

| **Frequencies (cm^-1^)** | **Assignment** |
| --- | --- |
| 3346 | O-H symmetric stretching |
| 2916 | Aromatic C-H stretching |
| 2849 | Aromatic C-H stretching |
| 1638 | C=C stretching |
| 1290 | C-O symmetric stretching |
| 1148 | S=O stretching |

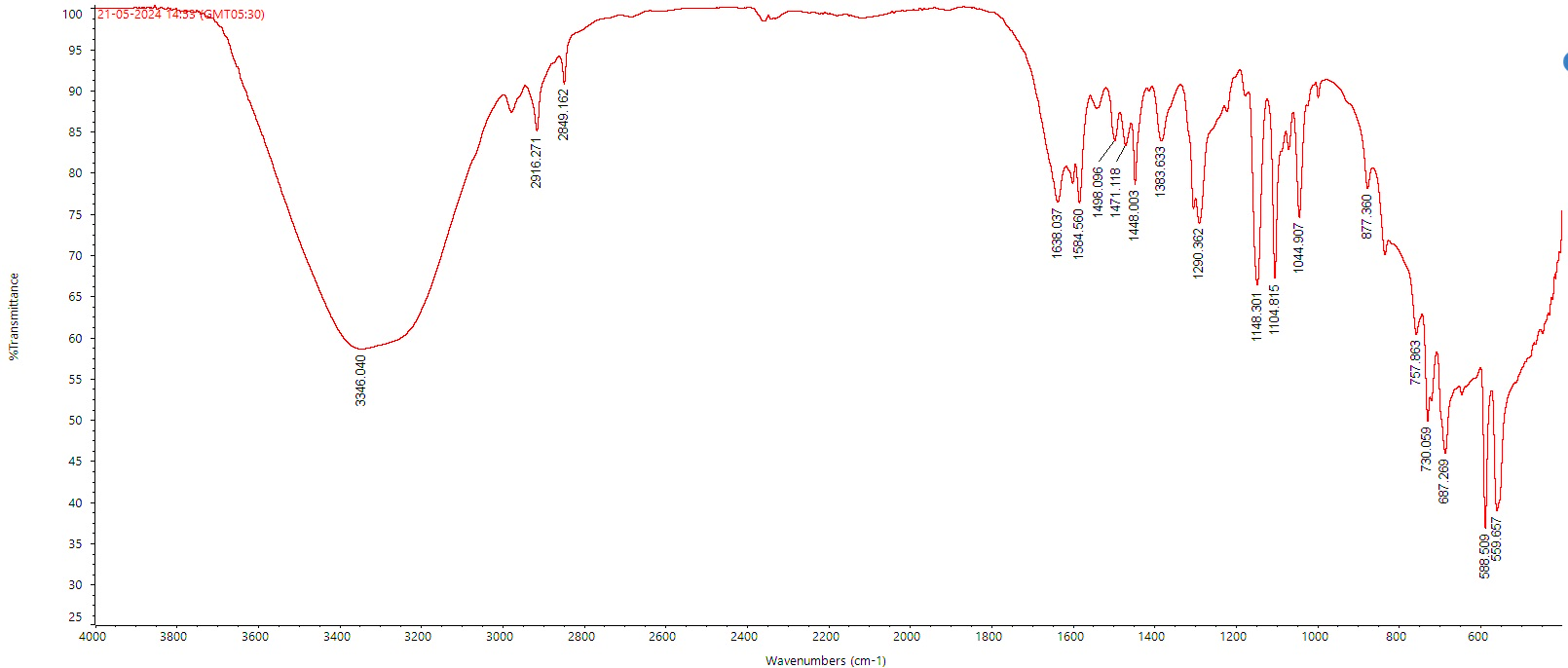

**Figure S1** FTIR spectrum of the chemicals extracted from the thermal bill paper

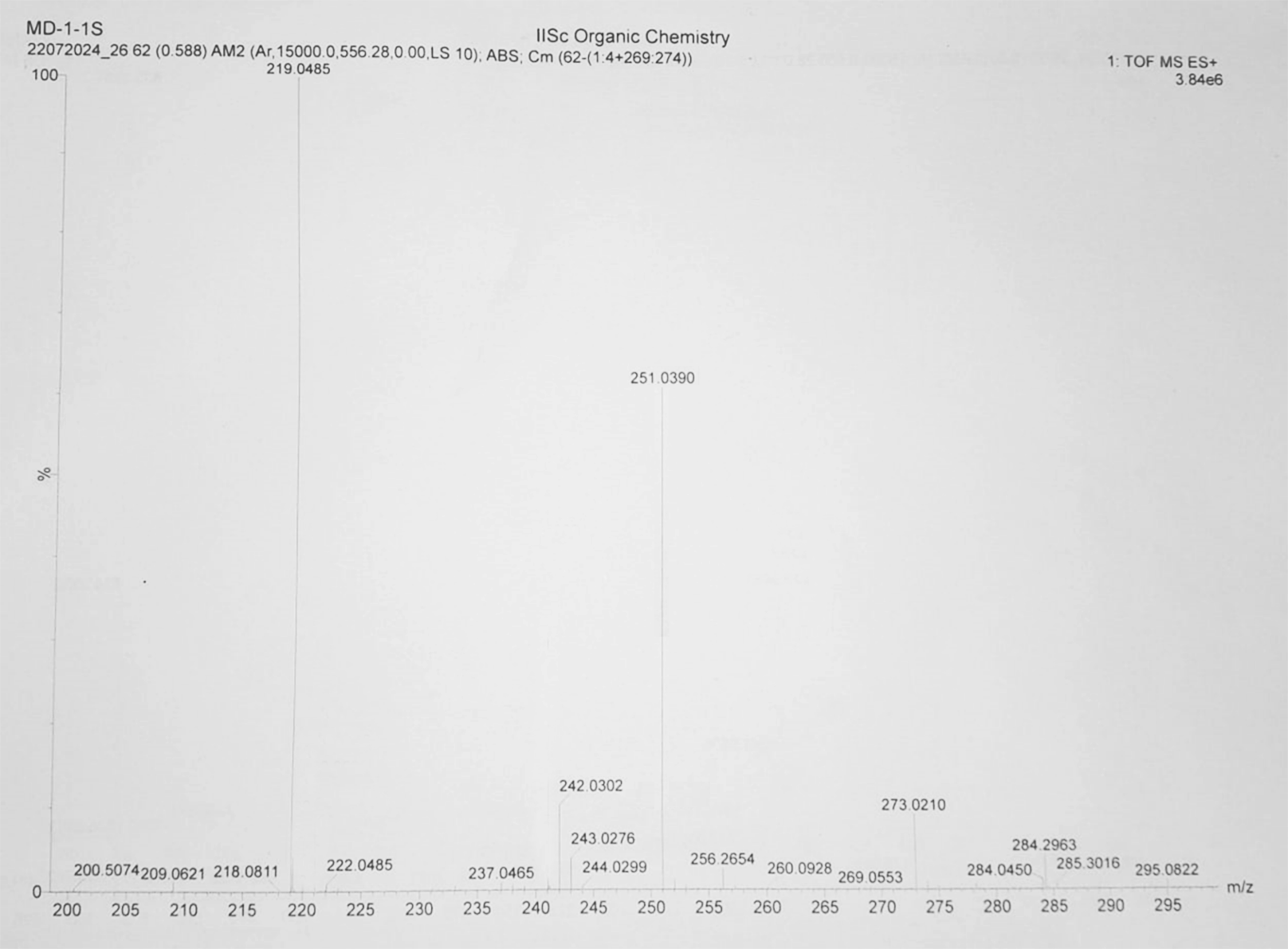

**Figure 14** High-resolution mass spectrum of the chemicals extracted from the thermal bill paper
